# Altered central pain processing in fibromyalgia – a multimodal neuroimaging case-control study using arterial spin labelling

**DOI:** 10.1101/2020.06.25.170647

**Authors:** Monika Müller, Florian Wüthrich, Andrea Federspiel, Roland Wiest, Niklaus Egloff, Stephan Reichenbach, Aristomenis Exadaktylos, Peter Jüni, Michele Curatolo, Sebastian Walther

## Abstract

Fibromyalgia is characterized by chronic pain and a striking discrepancy between objective signs of tissue damage and severity of pain. Function and structural alterations in brain areas involved in pain processing may explain this feature. Previous case-control studies in fibromyalgia focused on acute pain processing using experimentally-evoked pain paradigms. Yet, these studies do not allow conclusions about chronic, stimulus-independent pain. Resting-state cerebral blood flow (rsCBF) acquired by arterial spin labelling (ASL) may be a more accurate marker for chronic pain. The objective was to integrate four different functional and structural neuroimaging markers to evaluate the neural correlate of chronic, stimulus-independent pain using a resting-state paradigm. In line with the pathophysiological concept of enhanced central pain processing we hypothesized that rsCBF is increased in fibromyalgia in areas involved in processing of acute pain.

We performed an age matched case-control study of 32 female fibromyalgia patients and 32 pain-free controls and calculated group-differences in rsCBF, resting state functional connectivity, grey matter density and cortical thickness using whole-brain and region of interest analyses. We adjusted all analyses for depression and anxiety. As centrally acting drugs are likely to interfere with neuroimaging markers, we performed a subgroup analysis limited to patients not taking such drugs.

We found no differences between cases and control in rsCBF of the thalamus, the basal ganglia, the insula, the somatosensory cortex, the prefrontal cortex, the anterior cingulum and supplementary motor area as brain previously identified to be involved in acute processing in fibromyalgia. The results remained robust across all four neuroimaging markers and when limiting the study population to patients not taking centrally acting drugs and matched controls.

In conclusion, we found no evidence for functional or structural alterations in brain areas involved in pain processing in fibromyalgia that could reflect neural correlates of chronic stimulus-independent pain.

## Introduction

Fibromyalgia is characterized by chronic and widespread pain with additional symptoms such as fatigue, sleep disturbance and cognitive dysfunctions (1–3). The impact on quality of life is comparable to other chronic diseases such as rheumatoid arthritis, diabetes mellitus and chronic obstructive lung disease (4, 5). Therapeutic options remain limited with modest effects for most treatments and a high proportion of patients not responding to any treatment (6). Despite the clinical significance of the disease, pathophysiological processes remain poorly understood, which limits the development of diagnostic markers and novel treatments that target pathophysiological mechanisms rather than disease symptoms.

The striking discrepancy between objective signs of tissue damage and magnitude of pain suggests a pathophysiological process involving the central nervous system with possible alterations in brain function and structure (7, 8). A state of enhanced central pain response associated with increased neural activity in pain processing brain areas may lead to exaggerated pain response even to non-painful stimuli, high stimulus-independent pain and widespread pain (7, 9–12). Brain areas identified to be involved in pain processing include subcortical regions such as thalamus and basal ganglia and the insula, somatosensory cortex, prefrontal cortex, anterior cingulum and supplementary motor area as cortical regions (8, 13, 14). It remains however a subject of debate whether these brain areas are mainly processing acute pain signals or whether they are also involved in chronic pain (8, 15). Potential neuroimaging markers of chronic pain include resting-state cerebral blood flow (rsCBF) or functional connectivity on a functional level, as well as alterations of grey matter density and cortical thickness on a structural level.

Until now, neuroimaging studies investigating functional markers mainly focused on the comparison of acute pain processing mechanisms in fibromyalgia patients and pain-free controls using experimentally-evoked pain paradigms and blood oxygenation level dependent (BOLD) contrasts (7, 8). In line with the pathophysiological concept of enhanced central pain response, these studies detected increased neural activity in the amygdala, the insula, the somatosensory cortex and the cingulate cortex of fibromyalgia patients after painful stimuli. However, for two reasons, these studies do not allow conclusions about processing of chronic, stimulus-independent pain that typically remains constant over time (16–18). First, the studies used experimentally evoked acute pain paradigms rather than resting-state paradigms that are more likely to capture neural adaptation to chronic pain. Second, even if resting-state data were acquired, BOLD contrasts were used to quantify resting-state connectivity as marker of chronic pain. While BOLD provides a relative and indirect measure of brain activity, rCBF gives an absolute value that is directly related to local brain metabolism and thus appears perfectly suited to quantify neural activity in chronic pain.

Arterial spin labelling (ASL) as advanced neuroimaging method quantifies resting-state cerebral flow (rsCBF) as direct marker of neural activity at rest (15). First studies using ASL in different chronic pain conditions such as chronic low back pain (19, 20), postsurgical pain (21), trigeminal neuropathy (22), postherpetic neuralgia (23) and fibromyalgia (24) are emerging. Although these studies employed a resting-state paradigm, they only partly accounted for important confounders such as depression, age, and gender, or use of centrally acting drugs. This may explain why the results of the studies remain inconclusive. Additionally, previous studies did not integrate functional and structural neuroimaging by applying multimodal neuroimaging.

We conducted a multimodal neuroimaging study integrating functional and structural markers to explore the neural correlates of chronic pain in fibromyalgia. The primary objective was to compare rsCBF patterns of fibromyalgia patients and pain-free controls using ASL. In line with the paradigm of enhanced central pain response we hypothesized that rsCBF in areas involved in pain processing is increased in fibromyalgia. The secondary objective was to compare resting-state functional connectivity, grey matter density and cortical thickness between cases and controls assuming a reduction in these neuroimaging markers in fibromyalgia. We adjusted all our analyses for depression and anxiety and conducted a subgroup analysis limited to patients free of any centrally acting drugs.

## Material and methods

### Participants and study design

We performed a 1:1 frequency age-matched case-control study in fibromyalgia patients and pain-free controls using 10 years age bands. We centrally recruited right-handed (25) female participants at the University Hospital of Bern, Switzerland. We randomly sampled cases from a pool of 238 patients who were first diagnosed with fibromyalgia at either the Department of Rheumatology, the Department of Psychosomatic Medicine or the Pain Clinic between November 2013 and January 2015. We randomly sampled controls from a pool of 9253 women who presented during the same period at the Emergency Department. We stopped sampling when we reached the required sample size. We included cases if they were confirmed the diagnosis of fibromyalgia according to the diagnostic criteria of the American College of Rheumatology at repeat clinical examination at enrolment (2). We included controls if they did not suffer from any chronic pain disorder and were pain-free two weeks prior to neuroimaging. Common exclusion criteria for all participants were conditions interfering with the MRI acquisition; neurologic co-morbidity or history of neurosurgical intervention; psychiatric co-morbidity other than unipolar depressive disorder; end-stage somatic co-morbidity; intake of strong opioids or any psychopharmacological treatment other than anticonvulsants or antidepressants; inability to understand the consequences of study participation; and pregnancy. We performed the study according to a prospective protocol approved by the local ethics committee (KEK 43/13) and in accordance with the Declaration of Helsinki (26). All participants gave written informed consent.

### Assessment of socio-demographic, psychological and clinical characteristics

We assessed the following socio-demographic and psychological characteristics in all participants: age; education (higher education vs lower education); marital status (married vs not married); depression and anxiety. We considered participants with high school or university degree as having higher education. We used the Beck Depression Inventory version 2 (BDI-II) (27) and the State-Trait-Anxiety-Inventory (STAI) (28) to assess depression and anxiety in all participants. In cases we additionally assessed pain intensity, pain duration, spread of pain and disability. We used the Numerical Rating Scale to measure average pain intensity within the last 24 hours (NRS, 0 = no pain to 10 = worst pain) and the Fibromyalgia Impact Questionnaire to assess disability (FIQ, 0 = no disability, 100 = most severe disability) (29). We used the Widespread Pain Index to characterize the spread of pain with (WPI, 0 = no body region with pain to 19 = generalized pain affecting all body regions) (2). We recorded long-term daily intake of centrally acting drugs such as light opioids, antidepressants or anticonvulsants.

### Neuroimaging

#### Image acquisition

We performed multimodal neuroimaging at the Institute of Neuroradiology of the University Hospital of Bern to acquire four neuroimaging markers: resting-state cerebral blood flow (rsCBF); resting-state functional connectivity (rsFC); grey matter density (GMD); and cortical thickness (CT). We performed the MRI with a 3-Tesla Trio whole-body scanner using a 12-channel radio-frequency head coil (Siemens Medical, Germany). All study participants were instructed to lie quietly with eyes closed not thinking of anything particular during functional scans. We obtained the sequences in the following order.

First, high resolution anatomical T1* weighted images: 176 sagittal slices with 256 × 224 matrix points with a non-cubic field of view (FOV) of 256 mm × 224 mm, yielding a nominal isotropic resolution of 1 mm^3^ (i.e. 1mm × 1mm × 1mm), repetition time (TR) = 7.92ms, echo time (TE) = 2.48ms, flip angle = 16°, inversion with symmetric timing (inversion time 910ms).

Second, a set of 80 functional T2* weighted images using a pseudo-continuous arterial spin labelling sequence (pCASL) (30, 31): eighteen axial slices at a distance of 1.0 mm; slice thickness = 6.0 mm; FOV = 230 x 230 mm^2^; matrix size = 128 x 128, yielding a voxel-size of 1.8mm × 1.8mm × 6mm; TR = 4000ms; TE = 18ms. The gap between the labeling slab and the proximal slice was 90 mm; gradient-echo; echo-planar readout; ascending order; acquisition time 45 ms per slice. Slice-selective gradient 6 mT/m, post-labeling delay *w* = 1250 ms, tagging duration τ = 1600 ms.

Third, BOLD functional T2* weighted images were acquired with an echo planar imaging (EPI) sequence: 32 axial slices, FOV = 192×192 mm^2^, matrix size = 64×64, gap thickness = 0.75 mm, resulting in a voxel size of 3×3×3 mm^3^, TR/TE 1980ms/30ms, flip angle = 90°, bandwidth = 2232Hz/Px, echo spacing = 0.51ms, 460 volumes.

#### Selection of Region of Interests (ROIs)

Even though our main statistical approach was to perform whole-brain analyses, we also investigated several cortical Regions of Interests (ROIs) likely to be involved in pain processing in fibromyalgia. We a priori defined the following brain areas to be of interest based on a recent meta-analysis by Dehghan et al (8): Insula, left anterior and middle cingulate cortex (ACC, MCC), right Amygdala, superior temporal gyrus (STG), right lingual gyrus and left primary and secondary cortex (S I, S II). For functional analyses of rsCBF and rsFC we defined the exact coordinates of 11 ROIs according to this meta-analysis (8). We created a box around the coordinates with size = 27 voxels (216 mm^3^). For structural analyses we defined 10 ROIs based on the AAL-Atlas which provides volumetric regions for GMD analyses (32) and 23 ROIs based on the Destrieux-Atlas which is based on gyral and sulcal surface needed for reconstruction of CT (33).

#### Image preprocessing and calculation of functional and structural neuroimaging markers

T1* images were segmented into gray matter, white matter and cerebrospinal fluid, normalized to the Montreal Neurological Institute MNI space and smoothened using an 8 mm full-width at half maximum (FWHM) Gaussian kernel. We quantified resting-state cerebral blood flow (rsCBF, ml/100g/min) according to a previously applied, standardized protocol (34–36) using the following formula:

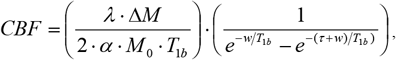

where *ΔM* is the difference between labeled and control image; *λ* the blood/tissue water partition coefficient (assumed 0.9); α the tagging efficiency (assumed 0.85); M_0_ the equilibrium brain tissue magnetization; τ the tagging duration; *w* the post-labeling delay; TI_2_ the image acquisition time; and T_1b_ the longitudinal relaxation time of blood (1650ms) (35). The resulting rsCBF maps were realigned and co-registered to the corresponding raw T1* image and normalized using the deformation matrix of the corresponding T1* image. We smoothed the resulting images with a 8mm FWHM Gaussian kernel. We then calculated grey matter rsCBF applying grey matter masks based on the segmented grey matter T1* images which were thresholded at 0.3. We conducted subject-wise first-level generalized linear models with rsCBF of the grey matter as outcome variable and the rsCBF of the white matter, the rsCBF of the cerebrospinal fluid and the realignment parameters as explanatory variables to correct for residual motion and artifacts. We checked six motion parameters (x-, y-, z-translations, roll, pitch, and yaw) and set a limit of two voxels of motion for exclusion. There was no significant difference between groups in motion and we did not exclude any subject due to excessive motion. We finally modelled a mean rsCBF map for each subject based on 40 pre-processed maps and computed global mean rsCBF within grey matter per subject based on this mean rsCBF map.

BOLD images were co-registered to the corresponding raw T1* image and then processed using the standard processing pipeline in CONN. This included realignment; slice-time correction; outlier-detection using ART-toolbox (global-signal z threshold 9, subject motion threshold 2 mm); normalization; smoothing with a 8 mm FWHM kernel; denoising by linear regression of white matter and cerebrospinal fluid signals, realigning and scrubbing parameters; and finally linear detrending as well as band-pass filtering between 0.008 and 0.09 Hz. Again, there was no significant difference between groups in motion and we did not exclude any subject due to excessive motion. We calculated rsFC between each pair of ROI for each subject by averaging and correlating the time-series of each ROI.

To compute GMD, we applied voxel-based morphometry to the pre-processed T1* images using standard processing modules with unmodulated segmentation in SPM12. To control for partial volume effects, we applied an absolute threshold of 0.2. We calculated the total intra-cranial volume by adding the segmented grey matter, white matter and cerebrospinal fluid volumes.

We carried out pre-processing and cortical reconstruction for CT analyses using the standard FreeSurfer package. This included automatic motion correction, segmentation, intensity normalization, inflation and registration to a spherical atlas.

### Statistical analysis

To explore differences in rsCBF, GMD and CT between patients and controls we performed uni- and multivariable generalized linear models based on whole-brain voxel-wise (vertex-wise for CT) analyses and ROI-analyses. The whole-brain analyses of rsCBF was the primary statistical analysis. We considered disease status (fibromyalgia patients vs pain-free controls) as explanatory variable and included rsCBF of total grey matter or total intracranial volume as co-variates in all analyses evaluating functional or structural neuroimaging markers, respectively. In adjusted analyses we additionally included the BDI-II and STAI-Trait t-value to consider co-morbid depression and anxiety. Age was corrected for in the study design using frequency matching of cases and controls according to age. To correct for multiple comparisons, we used family wise error (FWE) correction at peak-level in whole-brain and false discovery rate (FDR) in ROI-analyses. We set a cluster-threshold of 10 voxels. We explored group-differences in rsFC based on BOLD. We calculated crude and adjusted differences of rsFC for each pair of the 11 pre-defined ROIs. Again, we adjusted all analyses for BDI-ll and STAI-Trait t-values and applied FDR for multiple comparisons. We ran two sets of exploratory sensitivity analyses. First, we performed adjusted subgroup whole-brain analyses of the difference in rsCBF, GMD and CT between patients not taking any centrally acting drugs and their matched controls. Second, we explored the correlation of neuroimaging markers with clinical pain characteristics using Pearson correlation. We considered all neuroimaging markers with group-differences at p≤0.10 in any of the crude or adjusted analyses after correction for multiple comparisons for these correlations. We regarded correlation coefficients ≥0.7 as relevant. Due to the exploratory nature of these correlation analyses, multiple comparison correction was not employed. All reported p-values are two-sided and all confidence intervals refer to 95% boundaries. We performed statistical analyses in SPSS (Version 23.0. Armonk, NY: IBM Corp., ROI-analyses, correlations), SPM12 (Version 12, Welcome Trust, London, U.K.), Matlab (MATLAB 2015a; The MathWorks, Inc., Natick, MA, USA, whole-brain rsCBF, GMD), FreeSurfer (Version 5.3.0., http://surfer.nmr.mgh.harvard.edu/, CT) and CONN (Version 15, http://www.nitrc.org/projects/conn, rsFC).

## Results

### Study flow

We randomly selected and screened 151 fibromyalgia patients and 418 controls. The three most important reasons for excluding patients were neurologic or psychiatric co-morbidity other than unipolar depressive disorder (33 patients, 22%), inability to confirm fibromyalgia diagnosis at enrolment (23 patients, 15%) and inability to perform MRI (9 patients, 6%). The three most important reasons for excluding controls were neurologic or psychiatric co-morbidity (86 controls, 21%), chronic pain at enrolment (73 controls, 17%) and severe somatic co-morbidity (44 controls, 10%). We included 32 fibromyalgia patients and 32 age-matched pain-free controls. All participants were female and right-handed. S1 Fig presents the study flow diagram.

### Socio-demographic, psychological and clinical characteristics

Table 1 presents the socio-demographic and psychological characteristics of the study population. Cases and controls were comparable in terms of age, education, and marital status. Patients had significantly higher scores for depression (p≤ 0.001) and anxiety (p≤ 0.001). Twenty-four patients (75%) had widespread body pain affecting more than 50% of their body and 20 patients (63%) reported average pain intensity of at least NRS 6. All patients took pain medications on a daily basis; 21 (66%) of them regularly took centrally acting drugs such as weak opioids, antidepressants or anticonvulsants (intake of strong opioids was an exclusion criterion).

**Table 1.**
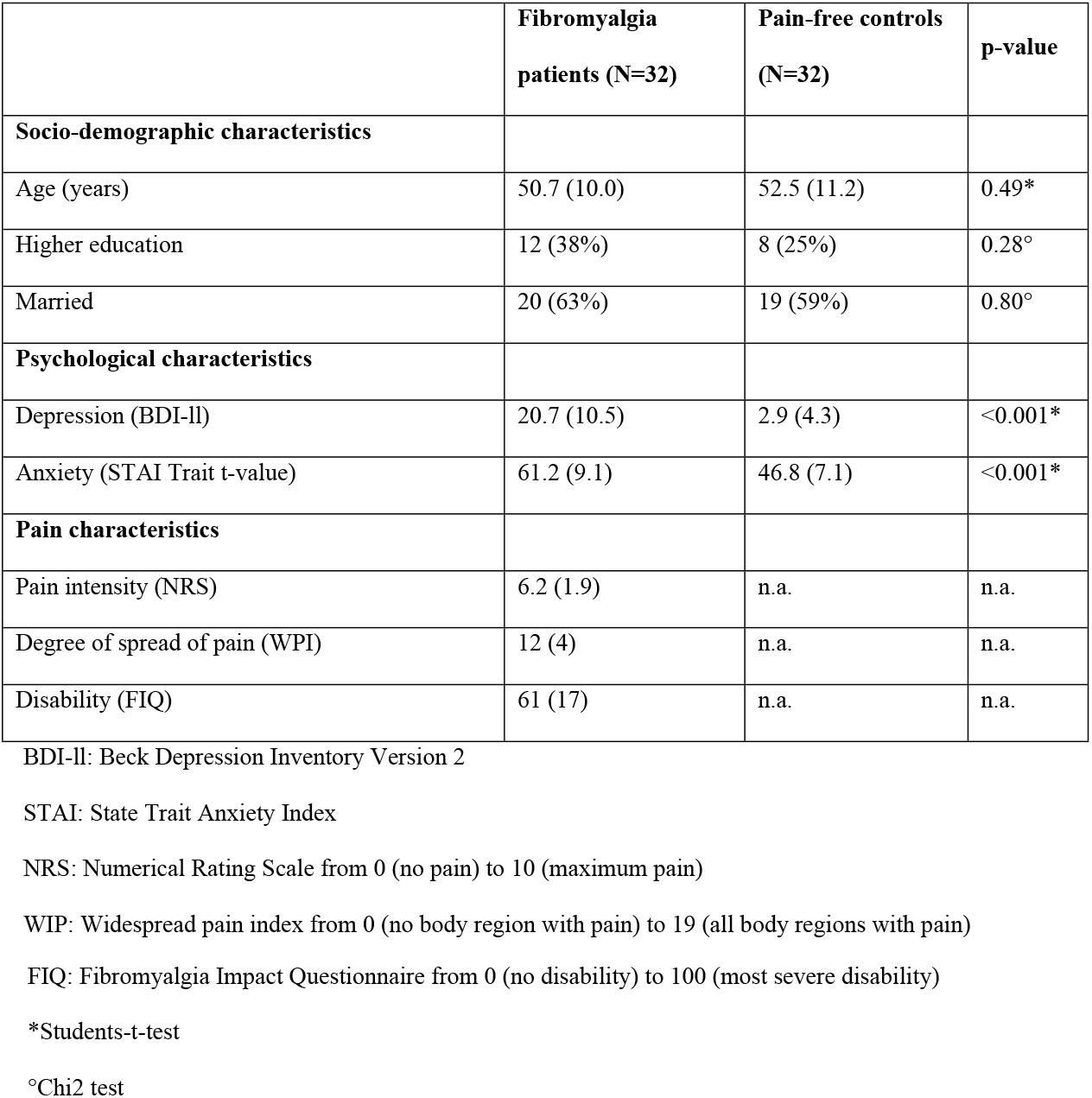
Baseline characteristics of fibromyalgia patients and pain-free controls. Values are numbers (percentage) or mean (standard deviation).

|  | **Fibromyalgia patients (N=32)** | **Pain-free controls (N=32)** | **p-value** |
| --- | --- | --- | --- |
| **Socio-demographic characteristics** |  |  |  |
| Age (years) | 50.7 (10.0) | 52.5 (11.2) | 0.49* |
| Higher education | 12 (38%) | 8 (25%) | 0.28° |
| Married | 20 (63%) | 19 (59%) | 0.80° |
| **Psychological characteristics** |  |  |  |
| Depression (BDI-II) | 20.7 (10.5) | 2.9 (4.3) | <0.001* |
| Anxiety (STAI Trait t-value) | 61.2 (9.1) | 46.8 (7.1) | <0.001* |
| **Pain characteristics** |  |  |  |
| Pain intensity (NRS) | 6.2 (1.9) | n.a. | n.a. |
| Degree of spread of pain (WPI) | 12 (4) | n.a. | n.a. |
| Disability (FIQ) | 61 (17) | n.a. | n.a. |

BDI-II: Beck Depression Inventory Version 2
STAI: State Trait Anxiety Index
NRS: Numerical Rating Scale from 0 (no pain) to 10 (maximum pain)
WIP: Widespread pain index from 0 (no body region with pain) to 19 (all body regions with pain)
FIQ: Fibromyalgia Impact Questionnaire from 0 (no disability) to 100 (most severe disability)
\*Students-t-test
°Chi2 test

### Differences in resting-state cerebral blood flow and correlation with pain characteristics

We found no increase in rsCBF in fibromyalgia patients as compared to pain-free controls in crude or adjusted whole-brain analyses. Contrary to our assumption, we found significant lower rsCBF in patients in the right Dorsolateral Prefrontal Cortex (x=44; y=32; z=26; cluster size =13 voxel; T= −4.39; p_FWE_=0.03) in crude analyses (Fig 1 A) and in the left Inferior Middle Temporal Gyrus (x=-50; y= −50; z=-4; cluster size= 110 voxel; T= −6.01, p_FWE_=0.002) in adjusted analyses (Fig 2 A). However, rsCBF in these two regions was not correlated with pain intensity, spread of pain or disability in 32 fibromyalgia patients (Fig 1 and 2B). Table 2 shows adjusted group-differences in rsCBF of 11 pre-defined ROIs. Patients showed numerically increased rsCBF in more than half of the areas (7 of 11 ROIs). None of the increases was statistically significant neither without nor with FDR correction. S1 Table shows crude differences of rsCBF of ROI-analyses. As in adjusted analyses, we were unable to detect relevant group-differences in crude analyses.

**Figure 1:**
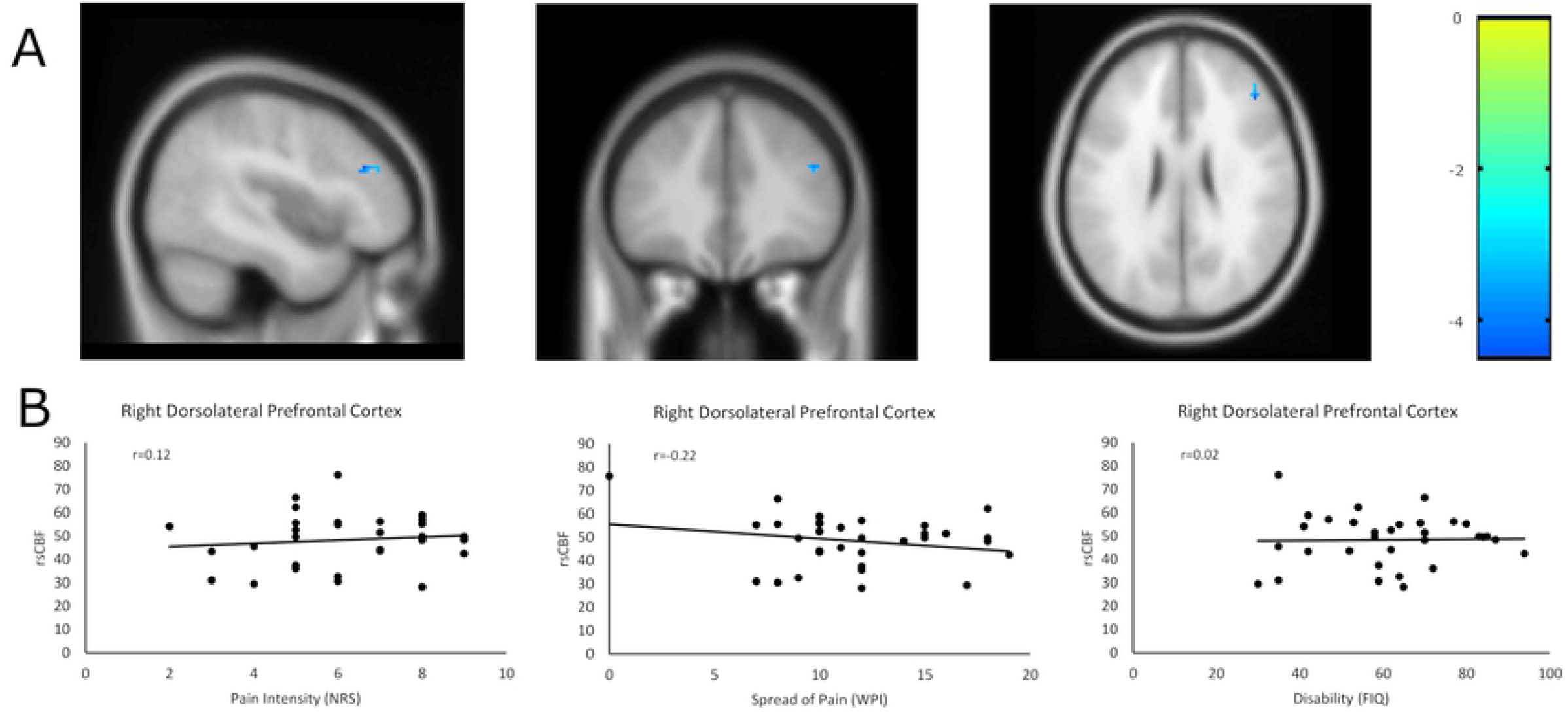
Crude difference in resting-state cerebral blood flow (rsCBF) between cases (N=32) and controls (N=32) in the right Dorsolateral Prefrontal Cortex (x=44; y=32; z=26) (A) and correlation of pain characteristics with rsCBF of this region within patients (N=32) (B). **A) Illustration of mean differences of rsCBF between groups. Colors indicate t-values controlled for global rsCBF.** **B) Scatter plots and correlation coefficients (r) of pain intensity; spread of pain; disability.** NRS: Numerical Rating Scale from 0 (no pain) to 10 (maximum pain) WIP: Widespread pain index from 0 (no body region with pain) to 19 (all body regions with pain) FIQ: Fibromyalgia Impact Questionnaire from 0 (no disability) to 100 (most severe disability)

**Figure 2:**
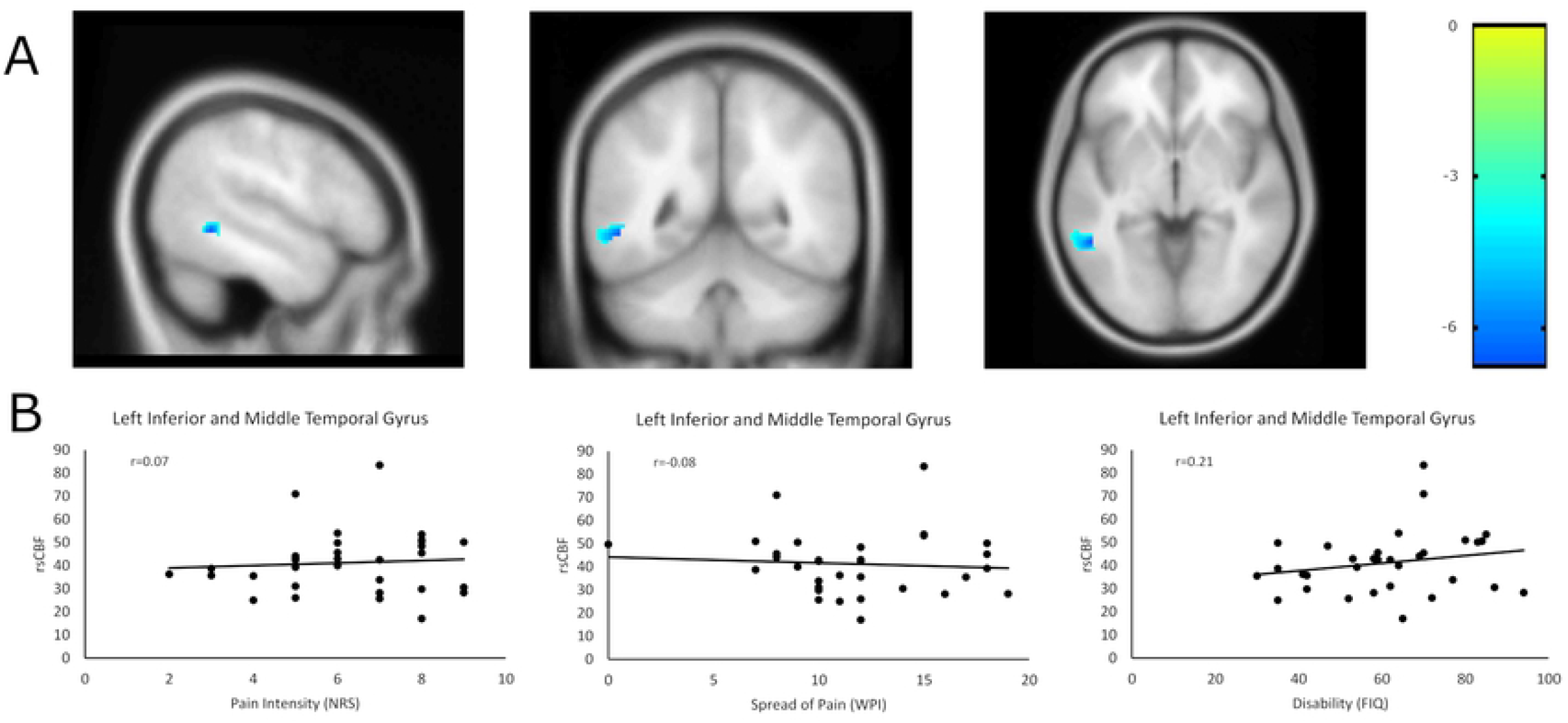
Adjusted difference in resting-state cerebral blood flow (rsCBF) between cases (N=32) and controls (N=32) in the left Inferior Middle Temporal Gyrus (x=-50; y= −50; z=-4) (A) and correlation of pain characteristics with rsCBF of this region within patients (N=32) (B). **A) Illustration of mean differences of rsCBF between groups. Colors indicate t-values controlled for global rsCBF and adjusted for depression and anxiety.** **B) Scatter plots and correlation coefficients (r) of pain intensity; spread of pain; disability.** NRS: Numerical Rating Scale from 0 (no pain) to 10 (maximum pain) WIP: Widespread pain index from 0 (no body region with pain) to 19 (all body regions with pain) FIQ: Fibromyalgia Impact Questionnaire from 0 (no disability) to 100 (most severe disability)

**Table 2.** Adjusted differences in resting state perfusion (rsCBF) between 32 fibromyalgia patients and 32 pain-free controls in 11 pre-specified Regions of Interest. Results are mean rsCBF with corresponding standard deviation and adjusted mean differences of rsCBF with corresponding 95% confidence intervals (CI), t-values and p-values from multivariable general linear models.

| MNI-coordinates |  |  | Brain area* | Mean rsCBF (SD) | Mean rsCBF (SD) | Mean difference | T-value | p-<br>uncorr |
| --- | --- | --- | --- | --- | --- | --- | --- | --- |
| x | y | z |  | fibromyalgia patients | pain-free controls | rsCBF (95% CI) |  |  |
| -48 | -27 | 24 | L insula | 36.65 (11.27) | 36.07 (11.12) | 1.97 (-5.98, 9.92) | 0.50 | 0.62 |
| 39 | 4 | 1 | R insula | 45.06 (13.63) | 43.43 (12.19) | 2.06 (-6.82, 10.93) | 0.46 | 0.65 |
| 43 | -2 | 1 | R insula | 45.74 (13.25) | 41.89 (14.70) | -1.57 (-11.46, 8.32) | -0.32 | 0.75 |
| -46 | -12 | 5 | L STG/insula | 52.60 (17.43) | 47.69 (14.60) | 1.24 (-10.34, 12.82) | 0.21 | 0.83 |
| 56 | -23 | 5 | R STG | 52.88 (12.50) | 50.46 (12.16) | 1.15 (-7.03, 9.33) | 0.28 | 0.78 |
| -15 | -48 | 72 | L SI | 25.52 (14.08) | 20.48 (45.91) | 10.15 (-16.50, 36.79) | 0.76 | 0.45 |
| -62 | -25 | 17 | L SII | 43.96 (11.01) | 42.49 (12.67) | 3.71 (-4.06, 11.47) | 0.96 | 0.34 |
| -2 | 48 | -12 | L ACC | 31.07 (19.80) | 35.97 (16.49) | -6.31 (-18.20, 5.59) | -1.06 | 0.29 |
| -17 | 30 | 31 | L MCC | 27.87 (10.07) | 27.39 (12.76) | -1.73 (-10.19, 6.74) | -0.41 | 0.68 |
| 13 | -55 | 7 | R lingual gyrus | 52.79 (13.08) | 47.92 (10.38) | 3.58 (-3.91, 11.07) | 0.96 | 0.34 |
| 27 | -12 | -15 | R amygdala | 47.79 (15.15) | 45.16 (13.26) | -4.14 (-14.63, 6.34) | -0.79 | 0.43 |

\* ROI selection based on meta-analysis by Dehghan et al. (Dehghan, M., et al., Coordinate-based (ALE) meta-analysis of brain activation in patients with fibromyalgia. Hum Brain Mapp, 2016. 37(5): p. 1749-58.)
Adjusted for rsCBF of grey matter, depression and anxiety MNI: Montreal Neurological Institute coordinates FDR: False Discovery Rate (correction for multiple comparison) L: Left, R: right STG: superior temporal gyrus, SII: secondary sensory cortex, SI: primary sensory cortex, ACC: anterior cingulate cortex, MCC: middle cingulate cortex

### Differences in resting-state functional connectivity, grey matter density and cortical thickness

We found no differences in rsFC among areas associated with chronic pain in fibromyalgia neither in crude not adjusted analyses after FDR correction. There was no evidence for differences in structural neuroimaging markers (GMD, CT) in patients as compared to controls in whole-brain analyses before or after FWE-correction. Table 3 and 4 report adjusted group-differences in GMD and CT of ROI-analyses. Patients showed non-significant decreases in structural neuroimaging markers in 8 of 10 ROIs for GMD and 12 of 23 ROIs for CT with and without FDR correction. S2 Table and S3 Table show crude differences of ROI-analyses of GMD and CT. As in adjusted analyses, we were unable to detect relevant group-differences in crude analyses.

**Table 3.** Adjusted differences in grey matter density between (GMD) 32 fibromyalgia patients and 32 pain-free controls in 10 pre-specified Regions of Interest. Results are mean GMD with corresponding standard deviation (SD) and adjusted mean differences of GMD with corresponding 95% confidence intervals (CI), t-values and p-values from multivariable general linear models.

| MNI coordinates |  |  | Brain area* | Mean GMD (SD) | Mean GMD (SD) | Mean difference GMD | T- | p- |
| --- | --- | --- | --- | --- | --- | --- | --- | --- |
| x | y | z |  | fibromyalgia patients | pain-free controls | (95% CI) | value | uncorr |
| -35 | 5 | 2 | L insula | 0.44 (0.05) | 0.45 (0.05) | -0.01 (-0.04, 0.02) | -0.63 | 0.53 |
| 39 | 5 | 1 | R insula | 0.43 (0.06) | 0.44 (0.05) | -0.01 (-0.04, 0.03) | -0.43 | 0.67 |
| -53 | -22 | 6 | L STG | 0.37 (0.04) | 0.38 (0.04) | -0.01 (-0.04, 0.02) | -0.56 | 0.58 |
| 58 | -23 | 5 | R STG | 0.35 (0.05) | 0.36 (0.04) | -0.01 (-0.04, 0.01) | -1.06 | 0.30 |
| -43 | -24 | 48 | L postcentral gyrus (SI) | 0.29 (0.03) | 0.30 (0.04) | -0.01 (-0.03, 0.02) | -0.58 | 0.56 |
| -47 | -10 | 13 | L rolandic operculum (SII) | 0.39 (0.05) | 0.40 (0.05) | -0.01 (-0.04, 0.02) | -0.59 | 0.56 |
| -4 | 34 | 13 | L ACC | 0.38 (0.05) | 0.39 (0.04) | 0.00 (-0.03, 0.03) | -0.20 | 0.84 |
| -6 | -16 | 40 | L MCC | 0.39 (0.05) | 0.40 (0.04) | 0.01 (-0.02, 0.03) | 0.43 | 0.67 |
| 16 | -68 | -5 | R lingual gyrus | 0.40 (0.04) | 0.42 (0.05) | -0.02 (-0.04, 0.01) | -1.26 | 0.21 |
| 27 | -1 | -19 | R amygdala | 0.51 (0.04) | 0.52 (0.04) | -0.02 (-0.04, 0.00) | -2.04 | 0.05 |

\* ROI selection based on AAL-Atlas (Tzourio-Mazoyer, N., et al., Automated anatomical labeling of activations in SPM using a macroscopic anatomical parcellation of the MNI MRI single-subject brain. Neuroimage, 2002. 15(1): p. 273-89.) Adjusted for total intracranial volume, depression and anxiety MNI: Montreal Neurological Institute coordinates. Coordinates are centres of mass.
FDR: False Discovery Rate (correction for multiple comparison)
L: left, R: right
STG: superior temporal gyrus, SII: secondary sensory cortex, ACC: anterior cingulate cortex, MCC: middle cingulate cortex

**Table 4.**
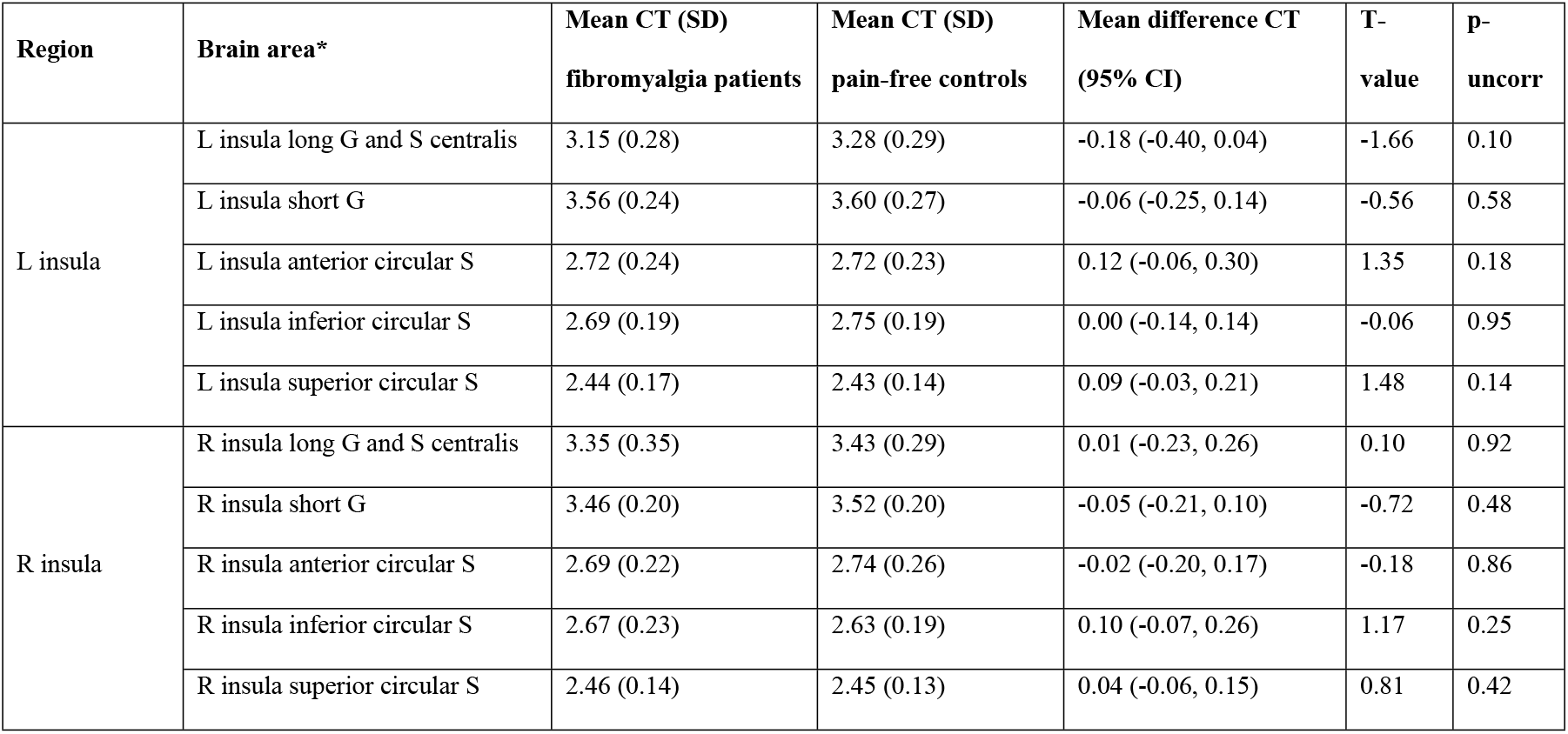
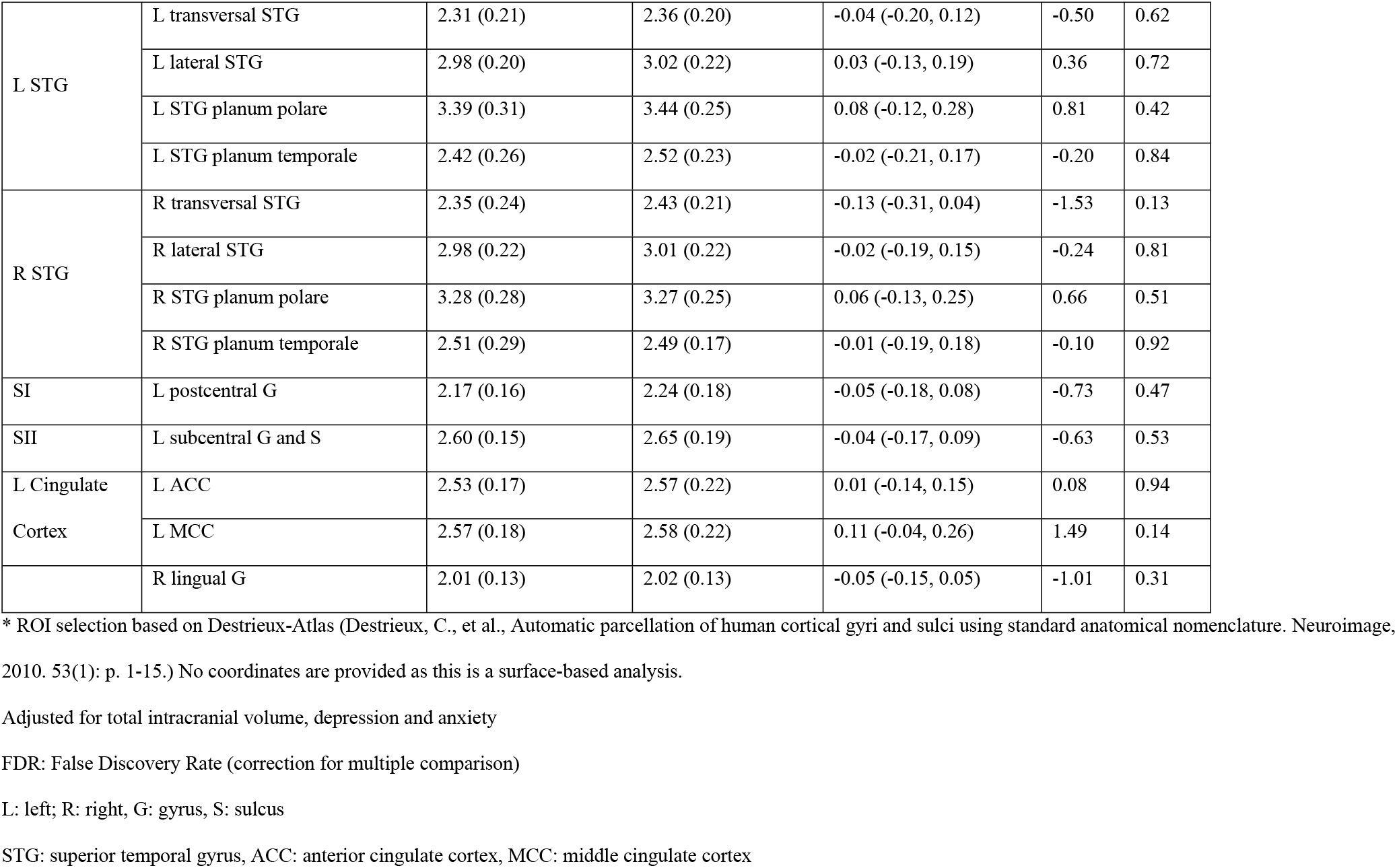
Adjusted differences in cortical thickness (CT) between 32 fibromyalgia patients and 32 pain-free controls in 23 pre-specified Regions of Interest. Results are mean CT with corresponding standard deviation (SD) and mean differences of CT with corresponding 95% confidence intervals (CI), t-values and p-values from multivariable general linear models.

### Subgroup analyses of patients not taking centrally acting drugs

Adjusted whole-brain subgroup-analyses of 11 patients not taking centrally acting drugs and matched controls showed no group-differences in rsCBF, GMD and CT.

## Discussion

### Main findings

To our knowledge, this is the first multimodal neuroimaging study integrating four different functional and structural markers of chronic pain in fibromyalgia. ASL was used to quantify rsCBF as measure of neural activity at rest and thus likely to reproduce a neuroimaging marker of chronic, stimulus-independent pain. Based on the pathophysiological concept of enhanced central pain processes, we expected an increased neural activity at rest in pain processing areas, i.e., an increased rsCBF in these brain areas in cases as compared to controls. Contrary to our hypothesis, we found no evidence of increases in rsCBF in brain areas involved in pain processing in fibromyalgia neither in whole-brain nor in ROI analyses. Instead, we found decreased rsCBF in patients in the right Dorsolateral Prefrontal Cortex in crude whole-brain analyses and in the left Inferior Middle Temporal Gyrus in whole-brain analyses adjusted for depression and anxiety, even after correcting for multiple comparisons. However, rsCBF in these two areas did not correlate with pain characteristics such as pain intensity, spread of pain or disability in patients. Additionally, and again contrary to our hypotheses, we did not find evidence for decreased rsFC, GMD or CT in brain areas involved in pain processing of fibromyalgia patients. The results remained robust in sensitivity analyses comparing fibromyalgia patients not taking centrally acting drugs with controls.

### Scientific context of our findings

There is ongoing debate to what extent brain areas processing acute pain signals are also involved in the development and maintenance of chronic, stimulus-independent pain (8, 15). We therefore defined the whole-brain analysis as the main statistical approach and performed secondary ROI analyses based on the recent meta-analysis by Dehghan and colleagues as most comprehensive evidence synthesis of functional and structural alterations in the central nervous system of fibromyalgia patients (8). Previous neuroimaging studies typically focused on the comparison of acute pain processing mechanisms between fibromyalgia patients and pain-free controls using experimentally-evoked pain paradigms (7). However, neuroimaging of chronic, stimulus-independent pain requires a different approach. Chronic pain typically remains constant during the course of an imaging session, rendering it invisible to traditional imaging techniques using pain paradigms. Task-free, resting-state parameters such as rsCBF are markers for brain activity at rest and thus more appropriate to measure chronic pain (16–18). ASL is the sequence of choice to measure rsCBF (15). Still, studies using rsCBF based on ASL are sparse in pain research with only one study performed to date in fibromyalgia (19–24). Although these studies employed a resting-state paradigm, none them integrated functional and structural neuroimaging by using multimodal scanning methods and none of them considered the effect of centrally acting drugs typically used by chronic pain patients on their results. Furthermore, only some of these studies addressed important confounders such as age, gender and concomitant depression.

To our knowledge, Shokouhi and colleagues conducted the only study using ASL in fibromyalgia (24). They compared 23 patients with 16 pain-free controls and thus included a much smaller and unmatched study population as compared to our study. They reported hypoperfusion in the putamen in subjects with chronic pain that was correlated with degree in disability but not pain intensity, thus suggesting that this hypoperfusion was due to adaptation processes. The correlation between rsCBF in the Putamen and disability was positive when using the Pain Disability Index to measure disability but negative when using the Fibromyalgia Impact Questionnaire as other measure of disability. The authors did not comment on this conflicting finding. However, they did not find group differences of rsCBF between fibromyalgia patients, which is in line with our findings. Previous studies investigating structural neuroimaging markers such as GMD or CT typically did not correct for co-morbid depression or for multiple comparisons, and if they did, group-differences in neuroimaging markers lost statistical significance (9–12). Therefore, our inability to find relevant group-differences in structural neuroimaging markers are in line with the findings of these previous studies (9–12).

### Strengths and limitations

Major strengths of the present study include: integration of four different functional and structural neuroimaging markers using a multimodal scanning protocol; evaluation of rsCBF using ASL; adjustment for depression and anxiety; correction for multiple comparisons and performance of a sensitivity analysis in a subgroup of patients not taking centrally acting drugs. Although fibromyalgia is characterized by intensive, widespread pain not associated with detectable peripheral lesions (7, 12), making it a good model to study the pathophysiology of enhanced central pain mechanisms, it also shows high co-morbidity with depression and anxiety. This may bias neuroimaging of chronic pain since brain areas involved in pain processing overlap with those involved in the pathophysiology of depression, e.g. prefrontal, cingulate, and supplementary motor cortex (34). Pain processing is tightly linked to top-down emotional control processes involving prefrontal cortices and amygdala (37). Hence, we adjusted all our analyses for depression and anxiety and found robust results.

Another problem of studies in fibromyalgia is the long-term treatment with centrally acting drugs, which may also interfere with neuroimaging markers. To avoid possible rebound-effects and for ethical reasons, we decided not to stop current medication and performed a subgroup-analysis including only patients not taking any centrally acting drugs and matched controls. We again found no group differences in whole-brain analyses of rsCBF, GMD and CT. We recruited both cases and controls in the only tertiary care hospital in the capital of Switzerland, with the same referral pathways for fibromyalgia patients and pain-free controls. This allowed us sampling cases and controls from the same source population, which is important to avoid selection bias in case-control studies.

Even though the present study is one of the largest studies in the field, the power to observe statistically significant group-differences may be limited with the consequence of possible false negative results. An argument for limited power to detect relevant group-differences is the fact that patients showed numerically increased rsCBF, decreased GMD and decreased CT in most of the pre-defined ROIs, even after adjusting for depression and anxiety, which would be in line with our hypotheses. Arguments that our inability to find group-differences was not merely due to a lack of power are the consistency of our results across four imaging modalities, the robustness to the type of analysis (whole-brain and ROI-analyses, sensitivity analyses) and the lack of correlation between neuroimaging markers and clinically relevant outcomes such as pain intensity, spread of pain and disability. Furthermore, by nature, case-control studies tend to inflate group-differences because severely sick cases are compared to healthy controls. This suggests that we would have detected moderate differences if they had been present.

### Implications

Our findings do not necessarily imply that altered central pain processing is not involved in fibromyalgia, but may point out that currently available functional and structural neuroimaging markers are not able to map stimulus-independent pain. Stimulus-independent, chronic pain may involve subtle functional and structural alterations which might be missed in medium-sized case-control studies like the present one and could be detected by larger multimodal studies also using resting-state designs. Furthermore, our study supports the hypothesis that brain areas processing acute pain signals are unlikely to be involved in the development and maintenance of chronic, stimulus-independent pain.

## Acknowledgments

We would like to thank Katrin Ziegler, MSc, Clinical Trials Unit, University of Bern, Switzerland, for her support in data-management and Fabienne Treichel, study nurse, for her support in patient recruitment and data collection. We owe our special gratitude to the Bangerter-Rhyner foundation for their financial support to this research project. No author has any conflicts of interest.

## Supporting information

**S1 Fig. Flow diagram of fibromyalgia patients and controls clinically evaluated between 1^st^ July 2011 and 30^th^ June 2013 and recruited for the study between 1^st^ November 2013 and 31^th^ January 2015 at the University Hospital of Bern.** $ according to criteria of American College of Rheumatology; *patients with light opioids, antidepressants, pregabalin or gabapentin included; °5 patients with mental retardation, 3 patients with pregnancy.

**S1 Table. Crude differences in resting state perfusion (rsCBF) between 32 fibromyalgia patients and 32 pain-free controls in 40 pre-specified Regions of Interest. Results are mean rsCBF with corresponding standard deviation (SD) and mean differences of rsCBF with corresponding 95% confidence intervals (CI), t-values and p-values from multivariable general linear models.**

**S2 Table. Crude differences in grey matter density between (GMD) 32 fibromyalgia patients and 32 pain-free controls in 10 pre-specified Regions of Interest. Results are mean grey matter density with corresponding standard deviation (SD) and mean differences of grey matter density with corresponding 95% confidence intervals (CI), t-values and p-values from multivariable general linear models.**

**S3 Table. Crude differences in cortical thickness (CT) between 32 fibromyalgia patients and 32 pain-free controls in 20 pre-specified Regions of Interest. Results are mean CT with corresponding standard deviation (SD) and mean differences of CT with corresponding 95% confidence intervals (CI), t-values and p-values from multivariable general linear models.**

